# Detection and assembly of novel sequence insertions using Linked-Read technology

**DOI:** 10.1101/551028

**Authors:** Dmitry Meleshko, Patrick Marks, Stephen Williams, Iman Hajirasouliha

## Abstract

**Motivation:** Emerging Linked-Read (aka read-cloud) technologies such as the 10x Genomics Chromium system have great potential for accurate detection and phasing of largescale human genome structural variations (SVs). By leveraging the *long-range* information encoded in Linked-Read sequencing, computational techniques are able to detect and characterize complex structural variations that are previously undetectable by short-read methods. However, there is no available Linked-Read method for detection and assembly of novel sequence insertions, DNA sequences present in a given sequenced sample but missing in the reference genome, without requiring whole genome de novo assembly. In this paper, we propose a novel integrated alignment-based and local-assembly-based algorithm, Novel-X, that effectively uses the barcode information encoded in Linked-Read sequencing datasets to improve detection of such events without the need of whole genome de novo assembly. We evaluated our method on two haploid human genomes, CHM1 and CHM13, sequenced on the 10x Genomics Chromium system. These genomes have been also characterized with high coverage PacBio long-reads recently. We also tested our method on NA12878, the wellknown HapMap CEPH diploid genome and the child genome in a Yoruba trio (NA19240) which was recently studied on multiple sequencing platforms. Detecting insertion events is very challenging using short reads and the only viable available solution is by long-read sequencing (e.g. PabBio or ONT). Our experiments, however, show that Novel-X finds many insertions that cannot be found by state of the art tools using short-read sequencing data but present in PacBio data. Since Linked-Read sequencing is significantly cheaper than long-read sequencing, our method using Linked-Reads enables routine large-scale screenings of sequenced genomes for novel sequence insertions.

**Availability:** Software is freely available at https://github.com/1dayac/novel_insertions

**Supplementary information:** Supplementary data are available at https://github.com/1dayac/novel_insertions_supplementary

## 1 Introduction

As a result of efforts in advancing DNA sequencing technologies and related algorithm developments for analyzing sequenced genomes, the field of personal genomics has been revolutionized in the past decade. Leveraging next-generation sequencing technologies, whole genome sequencing (WGS) has shown unprecedented promise in detecting and characterizing variants among human genomes as exemplified in the 1000 Genome Project (The 1000 Genomes Project Consortium (2010)). However, current methods using standard short-read sequencing are still unable to assemble a large fraction of structural variants due to limitations of short-reads in resolving repetitive regions of the genome effectively (Alkan *et al.* (2011); Chaisson *et al.* (2015b); Huddleston and Eichler (2016); Treangen (2012)).

Long-read sequencing technologies such as PacBio and Oxford Nanopore have recently become commercially available. These techniques promise the ability to resolve repetitive regions, call structural variants and improve de novo assembly (Chaisson *et al.* (2015a); Jain *et al.* (2018); Sedlazeck *et al.* (2017)). While these technologies offer much longer reads than traditional short-read technologies, their base-pair error rates are substantially higher than standard Illumina short reads (10-15% vs. 0.3% error) (Koren *et al.* (2012)). Additionally, long-read technologies have much higher costs (one to two orders of magnitude), lower throughput, and require large amounts of DNA as input (Lee *et al.* (2016); Rhoads and Au (2015)). This makes long-reads impractical for large-scale screenings of whole genome samples. The utility of current long-read technologies is therefore limited to a small number of targeted samples or validation purposes.

Low-cost, low-input and high-accurate Linked-Read technologies such as the 10x Genomics system have emerged recently to improve the ability of standard short-read sequencing technologies in determining whole genomes. In Linked-Read sequencing, DNA molecules are sheared into long fragments (10-100 kbp), and barcoded short reads from these long fragments are produced in such a way that reads from a long fragment share the same barcode. Such reads are referred to as *Linked-Reads* and the barcodes provide additional long-range information about a genome being sequenced. Most recently, Linked-Reads proved to be useful in multiple applications including but not limited to genome assembly (Bankevich and Pevzner (2016); Weisenfeld *et al.* (2017)), genome phasing (Kuleshov *et al.* (2014); Zheng *et al.* (2016)), metagenomics (Danko *et al.* (2019)), or large-scale somatic SV detection such as chromosomal rearrangements or general novel adjacencies (Greer *et al.* (2017); Spies *et al.* (2017)). The contribution of Linked-Reads to SV detection is, however, still limited to large structural variations (i.e. at least several thousand bp) and only certain classes of SVs. In particular, virtually none of the available SV detection algorithms attempt to characterize and assemble mid-size novel insertions i.e. DNA sequences as small as only 300 bp and up to a few thousand bp in size present in a given sequenced sample but missing in the reference genome.

In high-throughput sequencing experiments, the amount of generated reads determines the average sequence coverage of the target genome and is a direct indication of sequencing cost. With respect to sequence coverage, Linked-Read sequencing technologies consist of two key parameters, *C*_*F*_, and *C*_*R*_, which are defined as follows:

- *C*_*F*_ : The average coverage of the target genome with long fragments.
- *C*_*R*_: The average coverage of each long fragment with standard short reads.

Thus, the overall genome sequence coverage can be approximated as *C* = *C*_*F*_ × *C*_*R*_. A major goal in analyses of sequenced genomes is to keep the sequence coverage, and thus, the cost of sequencing as low as possible. In fact, with a relatively high *C*_*F*_ (e.g. 50x) and high *C*_*R*_ (e.g. 20x), we are able to assemble Linked-Reads and cover almost the entire target genome as was discussed in (Bishara *et al.* (2015); McCoy *et al.* (2014)) in the context of the Moleculo technology. However, this would require an enormous amount of sequencing. In the case of the latest 10x Genomics Chromium technology, which is the main focus of our study, *C*_*R*_ is very light. To obtain an average sequence coverage of 30*X*, the suggested parameters by the 10x Genomics system are *C*_*F*_ = 150*X* and *C*_*R*_ = 0.2*X*. Thus, it is impossible to assemble short reads sharing the same barcode. Therefore, in order to leverage the encoded barcode information for reference based or de novo assembly techniques, more sophisticated algorithms are needed.

Current techniques for detecting SVs using Linked-Reads mainly rely on quantifying barcode similarity of mapped reads between distant pairs of genomic locations to identify novel adjacencies in a target genome. In particular, Long Ranger (Marks *et al.* (2018)) and GROC-SVs (Spies *et al.* (2017)) state-of-the-arts SV discovery methods using Linked-Reads are very powerful in utilizing 10x long-range information to characterize regions with SV signatures that are at least 30 kbp apart. In this paper, however, we focus on detection of novel sequences insertions as small as 300 bp, a class of SVs that was not previously characterized by any existing Linked-Read method. While extremely challenging to characterize these insertions using short-read techniques because of the relatively short fragment lengths, such sequences may indeed contain functional elements and are of great interest as demonstrated by PacBio long-read analysis of such events (Huddleston *et al.* (2016)). Additionally, accurate identification of these insertions would help us build more platinum reference genomes and correct missassemblies in existing reference genomes.

In the past, several approaches attempted novel sequence insertion detection using standard short-read whole genome sequencing data. For example, NovelSeq (Hajirasouliha *et al.* (2010)) was the first algorithm developed to characterize these insertions using high coverage ultra short-read datasets (reads of length 35-41 bp). This algorithm was successfully applied to the 1000GP datasets and a number of NovelSeq calls were reported and validated (The 1000 Genomes Project Consortium (2010), Mills *et al.* (2011)). Subsequent short-read methods for this problem such as MindTheGap (Rizk *et al.* (2014)), ANISE and BASIL (Holtgrewe *et al.* (2015)), PopIns (Kehr *et al.* (2016)) or Pamir (Kavak *et al.* (2017)) appended population-based techniques (e.g. allowing multiple low-coverage samples from the same population) or used additional whole genome signals such as split-reads for better breakpoint resolution.

All algorithms above are based on the idea of assembling reads that are not aligned on the reference genome and connecting these assembled sequences with potential insertion breakpoints on the reference genome using paired-end information. NovelSeq (Hajirasouliha *et al.* (2010)) was the first algorithm capitalized on this idea. NovelSeq identifies unaligned pairedend reads with a single-end read aligned (i.e. One-End-Anchor reads) and performs a local assembly of those One-End-Anchor reads that clustered around same positions on the reference. Sequence contigs assembled in this way are simply called *anchors* and we use this term throughout this manuscript as well. An anchor represents a piece of sequence that can be located on the reference genome and can be used to find the breakpoint of a potential insertion event. NovelSeq then uses a de novo assembler such as ABySS (Simpson *et al.* (2009)) to assemble reads that none of their ends mapped to the reference (called orphan reads) and finally merges assembled contigs from the orphan reads with anchors using a greedy matching algorithm.

MindTheGap (Rizk *et al.* (2014)) uses a novel k-mer based signature to find insertion sites on the reference genome, while ANISE (Holtgrewe *et al.* (2015)) employs an elegant idea for resolving certain repeat copies. A more recent method, PopIns (Kehr *et al.* (2016)) presents an algorithm that uses information from different samples to find novel insertion sequences common to an ancestral population. The most recent algorithm Pamir (Kavak *et al.* (2017)) also generalizes NovelSeq’s approach for handling several low coverage genomes from the same population. Moreover, the accuracy and number of novel sequence insertions discovery were improved due to split-read signature usage and a non-greedy approach to match insertion sequences with their anchors. All these approaches are, however, limited to conventional paired-end sequencing data and it turns out to be problematic to correctly locate longer insertions (e.g. 300 bp or above), especially in the repetitive regions of the genome.

An explanation is that short-read libraries dramatically reduce our ability to locate an inserted sequence on the reference genome. Because the size of anchors is limited by the small insert size of the library and in most cases does not exceed the length of repetitive sequences common in mammalian genomes. Our main objective here is to develop a novel technique that can leverage barcodes and long fragment information encoded in Linked-Read sequencing to achieve much longer anchors. Such technique allows determination of the unambiguous location of novel sequence insertions on the reference even inside repetitive regions, a major limitation of short-read methods (see Figure 1 for a demonstration).

**Figure 1:**
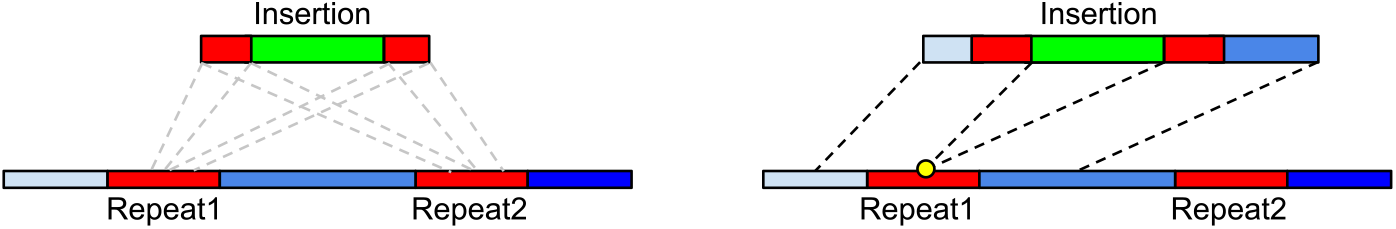
Schematic comparing novel insertion detection using standard short-reads with Linked-Reads. Shortcomings for existing short-read algorithms often arise from the fact that insertions can take place inside repetitive regions. Left: anchors found with paired-end data are too short to be uniquely placed on the reference genome as they can be mapped to any copy of repeat (shown in red). Right: Linked-Read sequencing data provides information that can be used for assembling long anchors that span repeat and the insertion breakpoint can be unambiguously placed on the reference genome.

Recently developed NUI-pipeline (Wong *et al.* (2018)) calls novel sequence insertions specifically with 10X data. It assembles whole dataset with SuperNova assembler and aligns poorly aligned reads into assembled contigs to identify insertion sequences and aligns contigs to the reference genome to find the position of breakpoints.

In what follows, we introduce an integrated mapping-based and assembly-based method, which is significantly more accurate than existing short-read methods for novel insertion discovery. While our method is less efficient that existing short-read methods, it is indeed more efficient compared to the recent Linked-Read algorithms that use whole-genome *de novo* assembly such as (Weisenfeld *et al.* (2017); Wong *et al.* (2018)) because it uses only a very small fraction of informative Linked-Reads as we describe below. While long-read sequencing is technically impractical for large-scale screening of whole genomes, our Linked-Read method is able to characterize one of the most challenging classes of SVs with a reasonable additional cost to standard short-read sequencing.

## 2 Methods

In this section, we describe our method, Novel-X, for detection of novel insertion sequences using Linked-Read sequencing. This method is based on a novel idea that the barcode information encoded in Linked-Reads can be used to reconstruct **long anchors** that can be unambiguously placed on the reference genome. This allows finding exact breakpoint positions on the reference even in certain repeat regions. Our approach is based on the local assembly of multiple barcodes originated from the same genomic loci.

The input for Novel-X is a reference genome (e.g. GRCh38) and a BAM-file produced by a read aligner, ideally, a 10x-specific aligner such as Lariat (Bishara *et al.* (2015)) available now in the Long Ranger package: (https://www.10xgenomics.com/software/). We refer to the set of the reads from the input BAM-file as *original reads* and to the BAM-file itself as *original BAM*. A pre-processing step in Novel-X is the extraction of paired-end reads from the original BAM that cannot be aligned to the reference genomes or have poor alignments. Intuitively, novel insertion sequences should consist of reads that do not align anywhere on the reference. The number of such insertions is not typically very high. Therefore, in contrast with de novo assembly based methods, we want to strictly filter out reads with high quality alignments to make the pipeline computationally effective. We choose paired-end reads in which at least one end is not aligned to the reference genome, or has the mapping quality below 10, or has more than 20% of soft-clipped bases. Additionally, they should have average phred-score above 20. For simplicity, we collectively call this set of unaligned reads as 𝒰. Reads from 𝒰 correspond to novel sequence insertions and anchor sequences.

In what follows, we present Novel-X via several steps.

1. **Novel sequence insertion assembly** – assembly of all reads from 𝒰. As the result of this step we obtain a set of novel insertion sequence candidates.
2. **Informative barcode list extraction** – for each insertion candidate we find barcodes with at least one read aligned to this insertion.
3. **Insertions reassembly** – we reassemble reads with barcodes found in the previous step to obtain long anchors for each insertion.
4. **Location of insertions on the reference** – we locate these anchors on the reference genome and find the exact position of each insertion.

We describe each step in details below. An overview of our technique is also shown in Figure 2.

**Figure 2:**
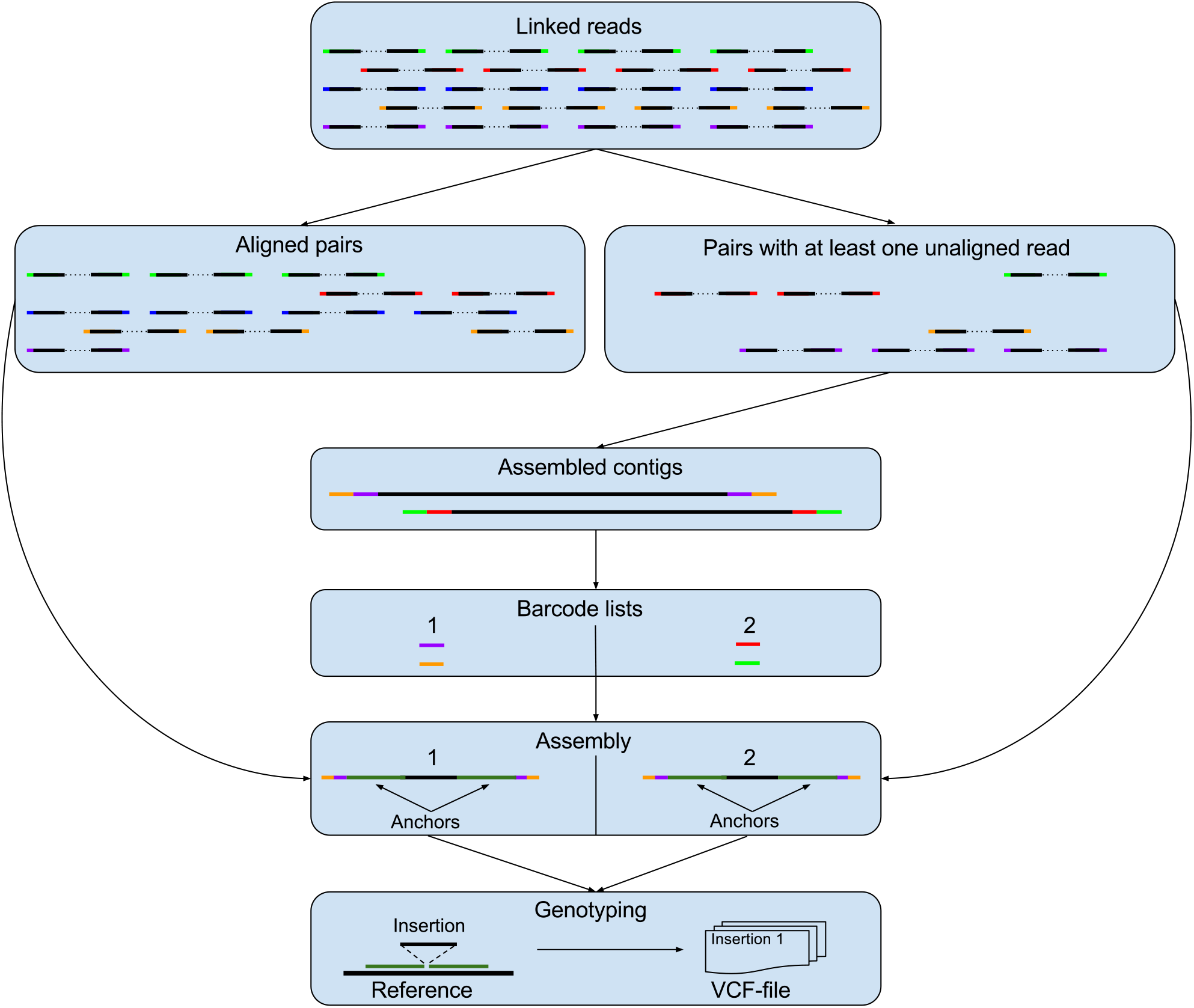
An overview of the steps in the Novel-X method is shown.

### 2.1 Novel sequence insertion assembly

We first use the Velvet *de novo* assembler (Zerbino (2010)) to assemble 𝒰. From our experience, Velvet is an assembler of choice for assembly of data with unusual properties. Other assemblers often have assumptions about the data such as diploidity or contiguity. We tried other assemblers at this step as well but results were shallow (in case of SuperNova) or highly chimeric (in case of SPAdes). Note that, *de novo* assembly of all reads in a high coverage whole genome sample is a computationally expensive task. However, 𝒰 consists only of a small fraction of the total reads and can be assembled efficiently. Ideally, the resulting assembly contigs would belong to sequences of novel insertions but could also be the results of misassembly or originate from contaminant sequences (i.e. non-human). If needed, we can perform a contamination removal procedure similar to what was previously done in NovelSeq (Hajirasouliha *et al.* (2010)) and Pamir (Kavak *et al.* (2017)). i.e. perform a BLAST search against nt/nr database and filter out all contigs that align to non-human references. Note that, we implemented this contamination removal procedure as an option in our software because it may also remove some known viral sequences (e.g. Human Herpesvirus sequence) that can be of interest for certain users. Similar to the analogy in (Hajirasouliha *et al.* (2010)), we call the remaining contigs *orphan contigs*.

### 2.2 Obtaining an informative barcode list

For each orphan contig *c*, we first align the reads from 𝒰 to *c* and filter read alignments with low-quality scores or with a large fraction (>20%) of soft-clipped or hard-clipped sequences. Note that, the exact definition of soft- and hard-clipped reads is aligner-specific. Intuitively, soft-clipped and hard-clipped read parts represent the part of a read that cannot be aligned together with the remainder of the read due to sequence dissimilarity. Let *R*(*c*) be the set of filtered barcoded-reads aligned to *c*. We denote *B*(*c*) as the set of all barcodes in *R*(*c*). We extract and store every read from the set of original reads whose barcode is in *B*(*c*). The information about barcodes of remaining reads is, however, extracted and aggregated separately for each orphan contig. Each contig that recruits less than *t* barcodes is discarded since the joint assembly of a limited number of barcodes is very unlikely to produce long anchors during the next step of the algorithm. The user-defined parameter *t* is set to 5 for a typical 10x whole human genome experiment by default.

### 2.3 Insertions reassembly

In order to reconstruct anchors and automatically connect them to novel sequences, for each barcode list we search the original BAM for reads that have a barcode from the barcode list and extract them. Then, we reassemble each set of extracted reads separately. By default, we use SPAdes (Bankevich *et al.* (2012)) with k=77 and coverage_cutoff=3, though usage of other options might be beneficial, e.g. using SuperNova is a good option if the sequence coverage is high. While we understand that different long fragments from different places can share identical barcodes, only regions of our interest would receive enough sequence coverage and can be assembled into contiguous sequences. Other sequences will be presented as extremely low-covered and thus should be filtered out during assembly (see Figure 3). Basically, all assembly pipelines have special procedures to process low-covered edges in the assembly graph. Usually, such edges in the assembly are considered as the result of sequencing errors or contamination in the data. While these edges might have different topological properties in the assembly graph, their negligible coverage will cause simplification procedures to delete them.

**Figure 3:**
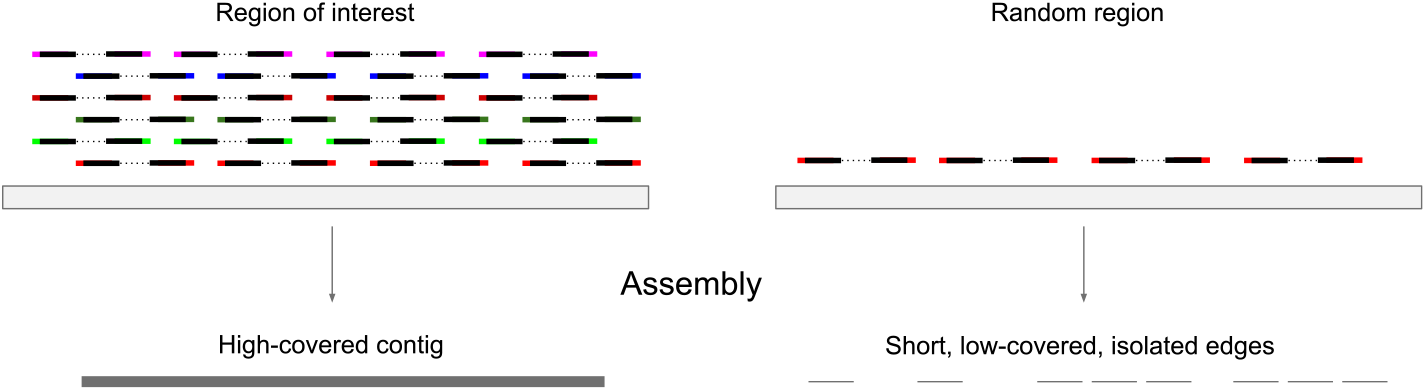
Underlying long fragment alignment to novel insertion site and some random genomic region and following assembly. Every barcode extracted from the barcode list recruits some underlying long molecule aligning to the insertion site. Long molecules with the same barcode can be recruited to align to the offsite genomic region but the probability of overlapping is close to 0 and assembly of such regions into contiguous sequences is unlikely.

In order to demonstrate that during this procedure random genomic regions would not be assembled into long contigs, we can model the genome as a set of non-intersecting regions and count the probability of aligning underlying long fragments in the same region. If the probability of getting a significant number of long fragments in a single bin is small, we can conclude that assembly of this region by chance is very unlikely. For example, if we divide the genome into bins of size 100 kbp (an approximate upper bound for 10X molecule length), in case of a haploid human genome we get approximately 30,000 bins. Empirically based on our experiments, the number of different barcodes that are recruited by a single novel sequence insertion of would be close to 50, because probability of getting of more that one read pair with the same barcode is small and read coverage in 10X experiments is close to 50. In that case, we roughly get 500 underlying long fragments that originated from a different genomic region assuming that on average 10 long fragments have the same barcode. Let us assume that one long fragment falls exactly to one bin. Given this simplification, we can find the probability of getting 0,1,2*, …* long fragments in a given bin combinatorially. We can then write a formula

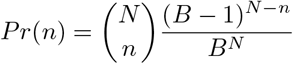

where *B* is a number of bins, *N* – number of long fragments and *n* – number of long fragments falling in a given bin. For the given values *Pr*(*n*) is equal to 0.9983, 0.0016, 1 × 10^-504^*, …* for *n* = 0,1,2, Moreover, each long fragment is only fractionally covered by short-reads (i.e. 0.1-0.2X), given that we do not expect any region with less than 5 barcodes recruited to be assembled. But if we were able to combine enough short reads from several long fragments originated from the same region, the assembly would be a feasible goal.

We also perform an additional filtering step using corresponding orphan contigs for each sample. We align orphan contigs to the set of contigs from the reassembly using Minimap2, the latest version of Minimap (Li (2016)). We perform the downstream analysis of only the contigs with the best match with orphan contig. Ideally, the result of this procedure can be presented as a single contig with “left anchor – insertion sequence – right anchor” structure.

### 2.4 Location of insertions on the reference

The last step of the pipeline is the detection of the positions on the reference genome where the novel insertions took place. We use Minimap2 for aligning the resulting assemblies to the reference genome. It allows us to align any number of candidate sequences to the human genome in a reasonable time with a high accuracy. Since Minimap2 results contain a large portion of short spurious alignments, we use a filtering procedure that resembles the QUAST (Gurevich *et al.* (2013)) procedure. i.e. choosing best alignment subsets from a given alignment set that maximizes the number of continuously covered basepairs. Remaining alignments that are adjacent with respect to the reference genome are analyzed for insertion signatures. We suppose that if the distance on the reference between adjacent alignments is small but large on the contig, then the contig contains an insertion and this insertion site is between these two alignments on the reference. The insertion content can be easily found as a subsequence on the contig between these alignments. For each insertion, we have right and left adjacent alignments that are considered as anchors for these insertions. We keep only insertions longer than 300 bp with at least one anchor exceeding 300 bp to prevent false calls. All found insertion are stored in a vcf (Variant Call Format) file.

## 3 Results

### 3.1 Benchmarking on simulated data

First, we evaluated Novel-X performance on data simulated from the hg38 reference genome. In order to do that, we inserted 124 sequences of 50-1000 bp length into the hg38 reference genome. To have a more realistic scenario, the list of sequences and positions was obtained from Huddleston and Eichler (2016) and resembles a list of non-template insertions in the CHM1 genome found with PacBio data. Reads were simulated using LRSIM (Luo *et al.* (2017)) and aligned back to the hg38 reference genome with LongRanger.

For all experiments and methods, we consider a pair of insertions as overlapping if their positions on the reference genome differ no more than 100bp. Among the 124 simulated insertion sequences, only 15 of them were of size more than 300bp. Novel-X correctly identified 14 of 15 insertions without any false positive calls. However, as we expect, Novel-X cannot assemble insertion sequences of smaller than 300 bp. Because, in order to successfully assemble a novel insertion, it needs a substantial number of barcodes to be associated with it. For very short insertions the number of aligned barcodes is usually small and the coverage is not sufficient for a successful assembly. For longer insertions, the number of barcodes is enough to assemble insertion with long anchors. Breakdown on the length of insertions is shown on Table 1

**Table 1:** Length breakdown of insertions from the simulated dataset. Novel-X found a majority of insertions longer than 300 bp.

| Novel insertions length breakdown for simulated dataset |  |  |  |  |
| --- | --- | --- | --- | --- |
| Length (bp) | 50-300 | 300-500 | $\geq 500$ | Total |
| Total | 109 | 7 | 8 | 124 |
| Novel-X | 6 | 6 | 8 | 20 |

### 3.2 Benchmarking on haploid hydatidiform moles

In order to evaluate the utility of Novel-X on real data, we performed experiments with a high coverage dataset generated from haploid genomes from two different hydatidiform moles (CHM1 and CHM13). Approximately 1ng of high molecular weight DNA was extracted and processed using a 10x Chromium instrument for each sample. The laboratory at 10x Genomics Inc. prepared these samples with the Chromium Genome reagents, Illumina-sequenced, and aligned to GRCh38. The datasets are available at https://support.10xgenomics.com/de-novo-assembly/datasets/2.0.0/chm.

Note that we chose these datasets because high coverage PacBio long-reads for these genomes were already publicly available. Furthermore, state-of-the-art short-read tools such as Pamir (Kavak *et al.* (2017)) for novel insertion detection used CHM1 for benchmarking. Unfortunately, Pamir is not designed to handle 10x sequencing data because the mrsFast aligner used inside Pamir’s pipeline requires paired reads with equal-length left and right reads. For a 10x data set, however, this assumption does not hold because the barcode sequence is actually encoded in the first read sequence. We tested Novel-X and Pamir on CHM1’s 10x dataset and WGS dataset both with high and sufficient coverage, and compared the results with the PacBio long-read data sets and results of Huddleston and Eichler (2016). Note that, we used 180 Gb of 10x data which is 56x. This would be equivalent to 130Gb Illumina short-reads at 56x given that the 10x barcode sequence takes up some of the read. Thus there is the need to sequence more on the 10x platform to compensate for sequencing the barcodes and achieve the same effective sequence coverage.

#### The CHM1 genome

In summary, for the CHM1 dataset, Novel-X identified 314 insertions longer than 300 bp with the mean length of 940 bp and a total length of 295 kbp. The average sum of the left and right anchors of insertion length equals 2,539 bp, while the standard deviation equals 2,444 bp. The maximum sum of two anchors length for a single insertion that we were able to achieve equals 15,821 bp. We also identified insertion sites that theoretically cannot be located on the reference genome with standard short-read data. Novel insertion detection tools for short-read data produce anchors that are limited by the insertion size (e.g. 300 bp for an anchor). Mapping such short sequences to repetitive regions is a hard and often a task that cannot be resolved unambiguously. In order to show that such insertion sites are ubiquitous, we extracted 300 bp upstream and downstream regions of insertion sites and searched these regions for repetitive sequences using RepeatMasker (Smit *et al.* (2004)). In total, for 77 insertions these regions consisted of repetitive sequences only. These results show that at least 25% of insertions found by our method would be hard to locate with WGS short-read data. Surprisingly, similar results were obtained for Pamir (27%) and PopIns (33%).

To compare results for different novel insertion callers we compared WGS novel sequence insertion callers with SMRT-SV data (see Table 2 and Figure 4). While Pamir finds more insertions than Novel-X and PopIns, it tends to call shorter insertions. Novel-X finds more insertion of size greater than 500 bp compared to other callers. These findings are consistent with our theory because longer insertion sequences recruit more barcodes than short ones and their assembly will more likely produce long anchors during the assembly step. Another encouraging validation of our method is that about 80% of Novel-X calls overlap with SMRTSV calls while the amount of agreement with Pamir and PopIns is below 45%. Indeed Pamir and PopIns are more likely to produce false positive calls. As it can be seen in Table 2, the number of PacBio calls is significantly higher than short-read methods’ calls because they are not necessarily just novel sequence insertions. For 15 novel sequence insertions longer than 300 bp reported in Chaisson *et al.* (2015a) using PacBio, Novel-X found 8, Pamir found 4 and PopIns found 2.

**Table 2:** Length breakdown and comparison between the PacBio based tool, SMRT-SV, short-read methods Pamir and PopIns, and our LinkedRead method Novel-X for CHM1. The numbers in brackets indicate the count of overlaps with SMRT-SV calls and the percentage of the overlapping calls. As it can be seen, the number of validated novel insertion calls with PacBio data using our method is significantly higher than those obtained by short-read methods.

| Novel insertions detection results for insertions longer than 300 bp |  |  |  |  |
| --- | --- | --- | --- | --- |
| Length (bp) | SMRT-SV | Novel-X | Pamir | PopIns |
| 300-499 | 1919 | 97 (78, **80%**) | 324 (**121**, 36%) | 156 (25, 16%) |
| 500-999 | 1144 | 139 (**115**, **83%**) | 76 (51, 67%) | 151 (33, 22%) |
| 1000-1999 | 598 | 56 (**49**, **88%**) | 5 (4, 80%) | 89 (13, 15%) |
| $\geq 2000$ | 608 | 22 (**12**, 55%) | 2 (2, **100%**) | 21 (6, 29%) |
| Total($\geq 300$) | 4269 | 314 (**254**, **81%**) | 407 (178, 44%) | 417 (77, 18%) |

**Figure 4:**
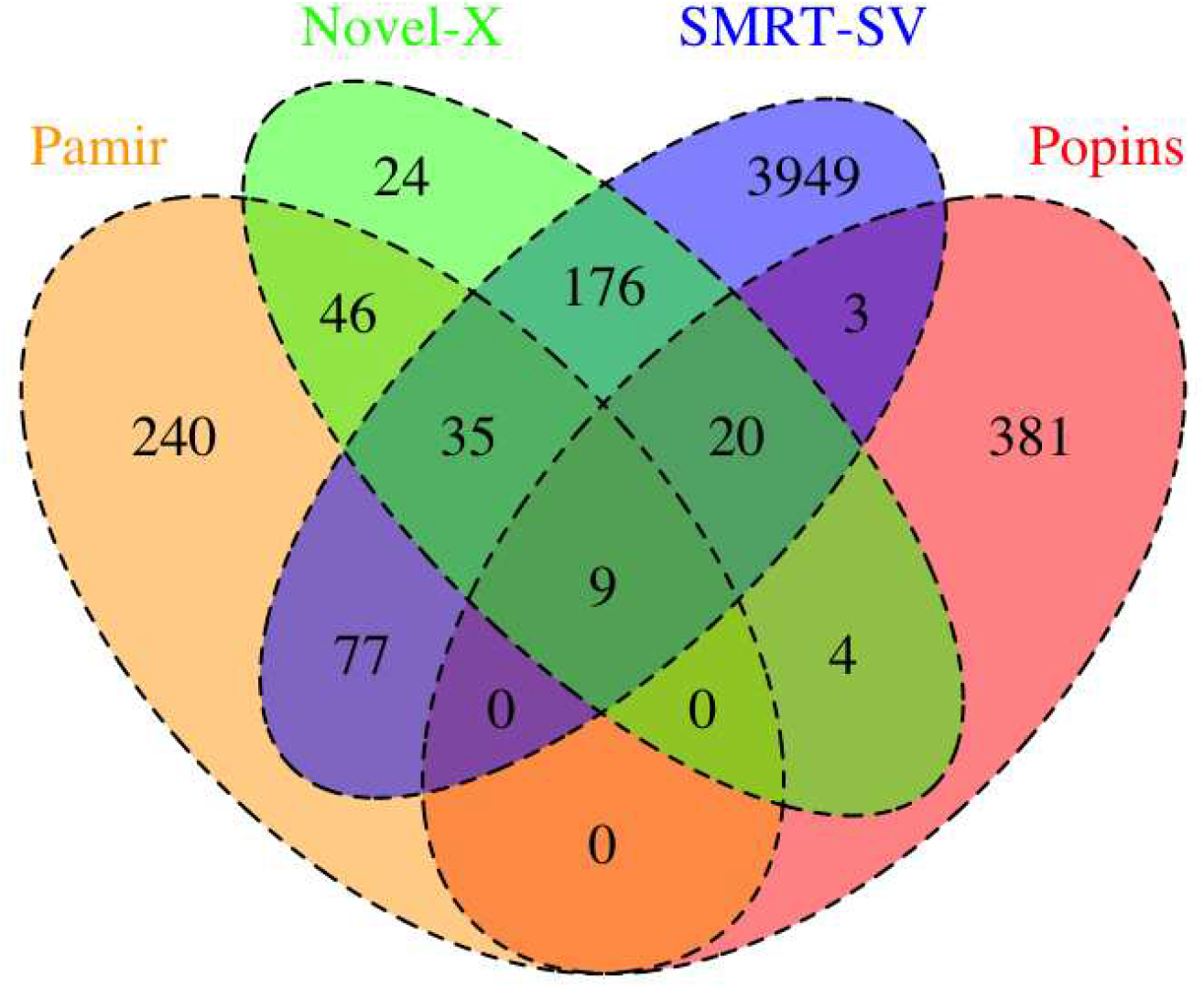
Euler-Venn diagram counting number of overlapping insertions between SMRT-SV, Novel-X, Pamir, and PopIns. Note that numbers for certain methods do not necessarily sum up to the values from Table 1. In certain cases, Pamir and PopIns provide multiple insertions calls with different insertion sequences but with the same location. Because CHM1 is a haploid genome, at most one call per location can be true. We add one to final numbers for each such event.

For insertions reported in Chaisson *et al.* (2015a), we also checked if sequence content for corresponding Novel-X/SMRT-SV and Pamir/SMRT-SV novel sequence insertions pairs is similar. In order to do that, we extracted insertion sequences for all methods and globally aligned overlapping insertion sequences maximizing number of matches and inspected identity scores and visual representation of alignments. For both methods for the majority of insertion pairs sequence percent identity tends to vary in 98-100% range, except few cases when insertion was truncated or extended. So for Novel-X we have two insertions extended for 31 and 20 nucleotides respectively and Pamir has insertion truncated by 226 nucleotides. This results shows that sequence content for all methods is similar for majority of the calls but ambiguities in calling step can have place.

#### Gene overlap analysis

We checked if novel insertion sequences overlap with known genes and coding sequences. In order to do that we compared breakpoint positions on the reference with gencode annotation v.24 (Harrow *et al.* (2012)). 150 out of 314 insertions falls inside known gene sequences, but only one of those falls into exon regions of known genes that are not pseudogenes. In general, it confirms our hypothesis that novel sequence insertions may contain novel exons or important non-coding regions but they don’t disrupt known exonic sequences.

Note that, while we focus on insertions longer than 300 bp in the main text, our method can indeed detect and report smaller insertions too (i.e. in the 50-300 bp range). See the Supplementary data available online where we provide information about all insertions found in CHM1. However, for such small insertions, we do not recommend Novel-X because we found that the long-range information used in our algorithm would not be helpful in this case. Given the current read length and short fragment sizes of standard Illumina sequencing, short-read techniques already have good performance for detecting these small events.

#### The CHM13 genome

We also applied Novel-X to the Linked-Read data set from the CHM13 sample. Novel-X found 293 novel sequence insertions with mean length 901 bp and a total length of 264 kbp. The average sum of the left and right anchors length equals 2,546 bp with standard deviation equals 2,136 bp, while the maximum sum of two anchors length for a single insertion that we were able to achieve equals 20,136 bp. Similar to the analysis of the CHM1 genomes, we also identified insertions that cannot be located on the reference genome with conventional short-read data. 72 out of 293 insertions fall into repetitive regions.

A large number of Novel-X calls overlaps with PacBio based SMRT-SV calls. For the CHM13 genome, out of 293 insertions longer that 300 bp found by Novel-X, 180 has a corresponding call in the PacBio dataset (see Table 3). Note that we did not run Pamir for that sample because of relatively large or infinite running times (more than 15 days with our hardware) but we expect the result to be in concordance with CHM1.

**Table 3:** Length breakdown and comparison between the PacBio based tool, SMRT-SV, short-read method PopIns, and our Linked-Read method Novel-X on CHM13. The numbers in brackets indicate the count of overlaps with SMRT-SV calls and the percentage of the overalpping calls.

| Novel insertions detection results for insertions longer than 300 bp |  |  |  |
| --- | --- | --- | --- |
| Length (bp) | SMRT-SV | Novel-X | PopIns |
| 300-499 | 1996 | 84 (**69**, **82%**) | 156 (37, 24%) |
| 500-999 | 1125 | 130 (**111**, **85%**) | 151 (40, 26%) |
| 1000-1999 | 591 | 53 (**36**, **68%**) | 89 (17, 19%) |
| $\geq 2000$ | 516 | 24 (**12**, **50%**) | 21 (6, 29%) |
| Total($\geq 300$) | 4228 | 291 (**228**, **78%**) | 417 (100, 24%) |

As an alternative way of validating our results, we compare the list of barcodes associated with every insertion with the list of barcodes for the region they were inserted. In order to do that, for each insertion we extracted barcodes in a 10 kbp window around its insertion site. We found that the barcode lists for majority of the insertions are completely of almost completely included in the corresponding barcode list for their insertion sites. This fact gives additional support to the hypothesis that our insertions are correctly placed on the reference genome (see Figure 5). We also counted barcodes aligned to 10 kbp window on the reference to barcodes associated with insertion ratios. We don’t expect these ratios to be high, because length of insertions is relatively small and they don’t recruit a lot of barcodes. The ratio values tend to vary between 0-20%, that meets our expectations.

**Figure 5:**
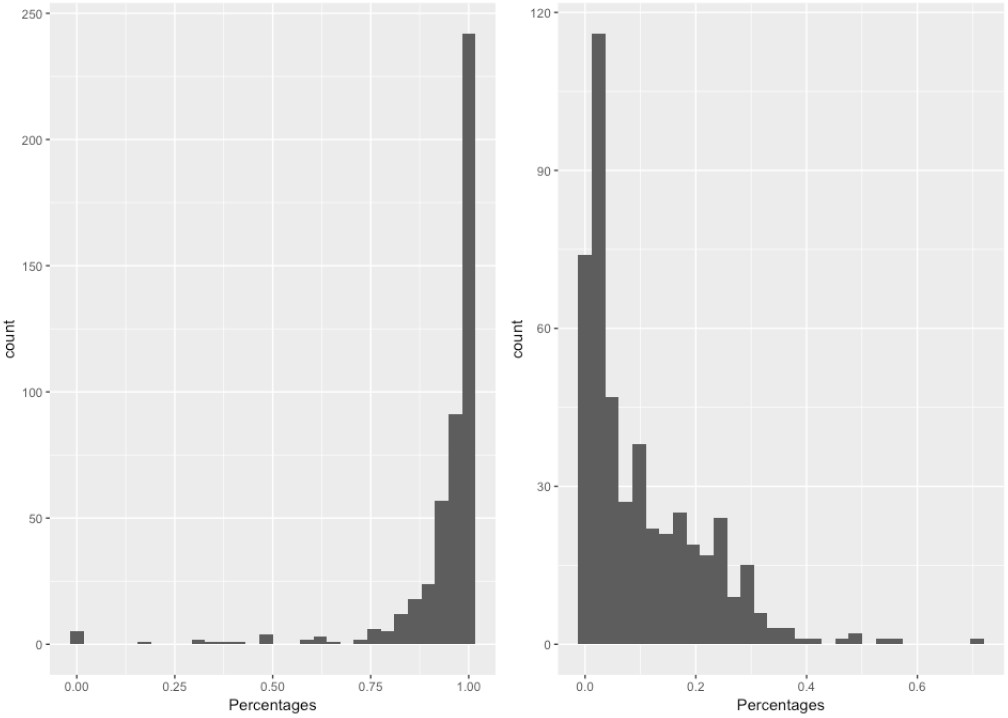
Left: Histogram of intersection of barcodes associated with insertions and found on the reference to associated with insertion barcodes ratio. Right: Histogram of barcodes aligned to 10 kbp window on the reference to barcodes associated with insertion ratios.

For novel insertions reported in Chaisson *et al.* (2015a), Novel-X called 6 insertions out of 12. We checked the sequence content similarity for corresponding Novel-X/SMRT-SV insertions the same way as for CHM1 dataset. For 5 insertions sequence similarity was in range 98-100%. Single insertion that had lower similarity was extended by 31 nucleotide comparing to SMRT-SV call.

### 3.3 The NA19240 Yoruba genome

We also tested Novel-X, Pamir and PopIns on a child from Yoruba trio from the 1000 Genomes Project The 1000 Genomes Project Consortium (2010) (NA19240). This genome was recently sequenced deeply on multiple platforms including 10x Chromium Chaisson *et al.* (2018). The coverage of this genome is 62x which makes it the deepest sequenced sample in our study.

In summary, for the NA19240 dataset Novel-X identified 478 insertions longer than 300 bp with the mean length of 1,123 bp and the total length of 536 kbp. Maximum insertion length found equals 29,906 bp. The average sum of the left and right anchors of insertion length equals 4,581 bp and the maximum sum of two anchors length for a single insertion that we were able to achieve equals 51,816 bp.

We compared insertions called with our method with SMRT-SV, NUI-pipeline, Pamir and PopIns calls. Results are summarized in Table 4. For this dataset, Linked-Read methods performed comparably to each other and better than their short-read counterparts. Novel-X is superior to NUI-pipeline in detecting and assembly of events more than 500bp, while the NUI-pipeline call set has more overlaps with the PacBio call sets for smaller insertions. Note that the NA19240 genome is sequenced at a higher coverage than typical sequencing experiments (i.e. 63x).

**Table 4:** Length breakdown and comparison between the PacBio based tool, SMRT-SV, short-read methods PopIns, Pamir, Linked-Read method NUI-pipeline and our Linked-Read method Novel-X on NA19240 data. The numbers in brackets indicate the count and percentage of overlaps with SMRT-SV calls.

| Novel insertions detection results for insertions longer than 300 bp |  |  |  |  |  |
| --- | --- | --- | --- | --- | --- |
| Length (bp) | SMRT-SV | Novel-X | Pamir | PopIns | NUI |
| 300-499 | 2453 | 168 (121, 72%) | 69 (56, 81%) | 232 (63, 27%) | 162 (**150**, **93%**) |
| 500-999 | 1183 | 189 (**155**, 82%) | 42 (39, **93%**) | 185 (42, 23%) | 66 (59, 89%) |
| 1000-1999 | 747 | 68 (46, 68%) | 14 (13, **93%**) | 76 (15, 20%) | 76 (**67**, 88%) |
| $\geq 2000$ | 743 | 53 (18, 34%) | 4 (4, **100%**) | 14 (2, 14%) | 77 (**65**, 84%) |
| Total($\geq 2000$) | 5126 | 478 (340, 71%) | 129 (112, 87%) | 507 (122, 24%) | 381 (**341**, **90%**) |

### 3.4 The NA12878 diploid genome

Finally, we ran our method on the well-known CEPH/HapMap NA12878 diploid genome (sequence coverage 52x) and compared it with a recently de novo assembly based method Wong *et al.* (2018) which we will refer to as NUI-pipeline). We obtained high-coverage 10x Chromium Linked-Reads from the publicly available Genome In A Bottle (GIAB) data set (Zook *et al.* (2018)). 1.25 billion paired 98 bp reads from the GM12878 cell line were extracted using the Chromium kit and aligned on the GRCh38 reference genome using Long Ranger 2.2 software.

For the NA12878 sample, Novel-X found 219 novel sequence insertions with mean length 778 bp and a total length of 170 kbp. The average sum of the left and right anchors length equals 4,404 bp with standard deviation equals 4057 bp, while maximum sum of two anchors length for a single insertion that we were able to achieve equals 28,247 bp. Analysis of the insertions that fall into repetitive regions was performed: 36 out of 219 insertions fall into repetitive regions.

We compared Novel-X calls with remapped (from hg37 to hg38 reference genome) SMRTSV PacBio call set from the GIAB project and NUI-pipeline calls.

PacBio dataset consists of 6,423 insertions of length more than 300 bp. Novel-X has 149 calls overlapping with this dataset. These results show that Novel-X has consistently high overlap ratio with PacBio data and can work in diploid settings.

For NA12878 sample we also tested recently published NUI-pipeline (Wong *et al.* (2018)). Surprisingly, it identified only 31 insertions longer than 300 bp (37 insertion of any length in total) and almost all of them overlap with PacBio calls (28 of 31). However, sensitivity is very low comparing to any other short-read method. Also, NUI-pipeline requires whole genome assembly and has a higher computational cost compared to short-read methods and Novel-X. Table 5 summarizes these results. These results suggest that our method, Novel-X performs much better than the competitor tool, NUI-pipeline, on the NA12878 with the sequence coverage of 52x (i.e. 20% lower than the sequence coverage of NA19240 in our study).

**Table 5:**
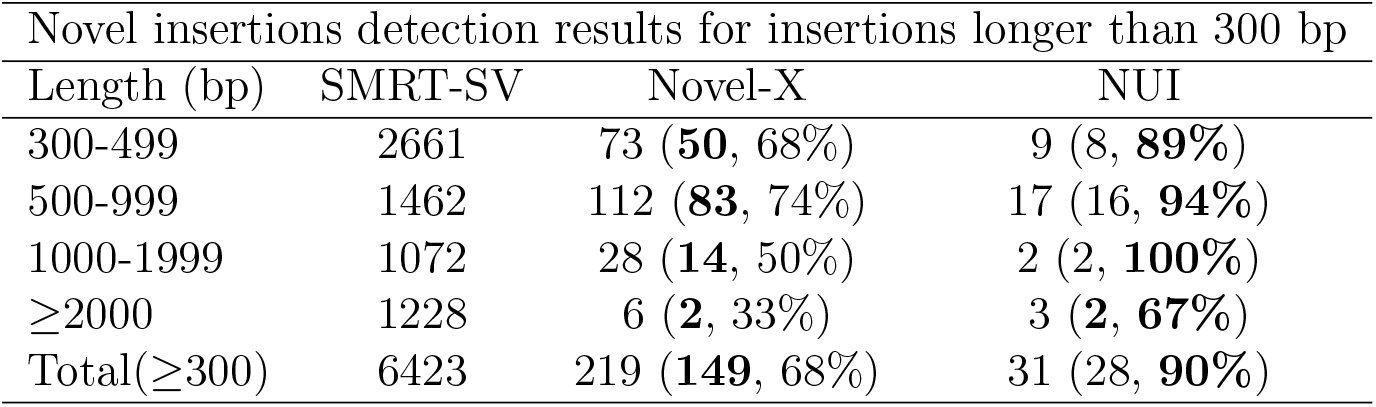
Length breakdown and comparison between the PacBio based tool, SMRT-SV, Linked-Read method NUI-pipeline and our Linked-Read method Novel-X on NA12878. The numbers in brackets indicate the count of overlaps with SMRT-SV calls.

| Novel insertions detection results for insertions longer than 300 bp |  |  |  |
| --- | --- | --- | --- |
| Length (bp) | SMRT-SV | Novel-X | NUI |
| 300-499 | 2661 | 73 (**50**, 68%) | 9 (8, **89%**) |
| 500-999 | 1462 | 112 (**83**, 74%) | 17 (16, **94%**) |
| 1000-1999 | 1072 | 28 (**14**, 50%) | 2 (2, **100%**) |
| $\geq 2000$ | 1228 | 6 (**2**, 33%) | 3 (**2**, **67%**) |
| Total($\geq 300$) | 6423 | 219 (**149**, 68%) | 31 (28, **90%**) |

### 4 Discussion

In this paper, we described a novel strategy for local assembly of multiple barcodes originated from the same genomic loci. We believe as a future work, this strategy has the potential to be used in the assembly of other classes of complex SVs. Such strategy greatly decreases the complexity of genomic loci assembly even with a complex repeat composition because these repeats can be distinguished using the encoded barcode information.

While the Novel-X approach runs slower that existing short-read methods (e.g. PopIns), it is able to found more insertions confirmed with PacBio data. Another alternative is to use whole genome assembly (e.g. NUI-pipeline). However, whole genome assembly requires more computational resources than Novel-X. Typical peak memory usage for Novel-X is around 100Gb (during assembly of 𝒰) and for SuperNova 2.1 whole genome assembly it goes up to 250Gb. In order to achieve the same 3-day time consumption, Supernova assembly requires 26 cores, while typical Novel-X run requires only 8. So, Novel-X is more accurate than short-read methods and more effective while maintaining comparable accuracy that whole genome assembly methods. It makes Novel-X a reasonable method for novel sequence insertion calling.

One of the problems we faced during our method development was the choice of an assembler. Most of the algorithms that use assembly-based techniques use Velvet (Zerbino (2010)) as the assembler of choice. We also used Velvet for this study because it is a reliable and conservative choice. Note that Supernova 2.0, a newer release of the original Supernova software developed by a team at the 10x Genomics Weisenfeld *et al.* (2017) can be also an alternative choice. In the case of SuperNova assembler, however, the user has no control for any assembler parameters and default parameters are maximally tuned for whole human genome diploid assembly. From our experience with SuperNova with default parameters, we believe that some viable novel insertion candidates were dropped out during the assembly phase due to low coverage. A future direction would be to develop a specific local assembler designed for SV detection tasks in Linked-Read data.

While we mainly validated our call sets using orthogonal long read technologies, a future work would be to use PCR and Sanger sequencing to further validate our Novel-X predictions, especially for those calls that were not found in the PacBio dataset. Furthermore, while other long-read techniques such as Oxford Nanopore (ONT) become more and more available, there will be alternative opportunities to validate calls sets produced by our method.

## Acknowledgements

We sincerely thank Yin-Yi Lin, John Huddleston, Mark Chaisson, and Evan Eichler for providing assistant with Pamir and SMRT-SV call sets. We acknowledge the laboratory in 10x Genomics Inc. for library preparation and sequencing the Linked-Read data used in this study. We also thank David C. Danko for helpful discussions and proofreading the manuscript.

